# Extrachromosomal DNA in cutaneous squamous cell carcinoma is associated with increased nodal disease

**DOI:** 10.1101/2024.02.04.578845

**Authors:** Tomas Bencomo, Amarinder Thind, Bruce Ashford, Marie Ranson, Carolyn S. Lee

## Abstract

Extrachromosomal DNA (ecDNA) is a mechanism of oncogene amplification that occurs in many cancers and has been linked to worse patient outcomes. While ecDNA has previously been identified in cutaneous melanoma, it has yet to be described in cutaneous squamous cell carcinoma (cSCC). Using whole genome sequencing and transcriptomic data of primary and metastatic cSCC tumors, we establish the presence of ecDNA in cSCC and confirm elevated expression of ecDNA-associated genes. We also find that the presence of ecDNA correlates with a higher lymph node ratio, a measure of disease severity. Together, these findings are the first to report ecDNA in cSCC and suggest that ecDNA gene amplifications promote an aggressive cSCC phenotype.

## Main Text

Extrachromosomal DNA (ecDNA) is a mechanism of oncogene and regulatory element amplification in many cancers that is often associated with worse clinical outcomes (Kim et al. 2020). In contrast to chromosomes that are distributed equally to daughter cells during cell division, ecDNA segregates randomly during mitosis, accelerating tumor evolution and giving rise to genetic heterogeneity that may underlie drug resistance and tumor aggressiveness (Lange et al. 2022). While extrachromosomal amplifications may underlie treatment non-response in melanoma (Spain et al. 2023), characterization of ecDNA in other cutaneous malignancies remains undefined. Here we establish the frequency and characteristics of ecDNA in primary and metastatic cutaneous squamous cell carcinomas (cSCC) from 40 patients and find that the presence of ecDNA is associated with a higher lymph node ratio, suggesting it may indicate more extensive disease.

AmpliconArchitect (Turner et al. 2017) detected ecDNA in 21% (5 of 24) of cSCC metastases analyzed by whole-genome sequencing (Thind et al. 2022), but never in matched normal blood (0 of 24) (**Table 1**). By comparison, ecDNA was identified in 11% (2 of 19) of primary cSCC (Gupta et al. 2023), with no significant association found between ecDNA presence and primary vs. metastatic status (Fisher exact test, P=0.437). After accounting for three patients that provided both primary tumor and metastasis, the overall detection frequency of ecDNA in cSCC was 15% (6 of 40 unique tumors). These findings confirm the absence of ecDNA in normal tissue and demonstrate that the ecDNA amplification frequency in cSCC is comparable to that reported across 25 cancer types (14%) including closely related head and neck cancers (25%) (Kim et al. 2020).

**Table 1.**
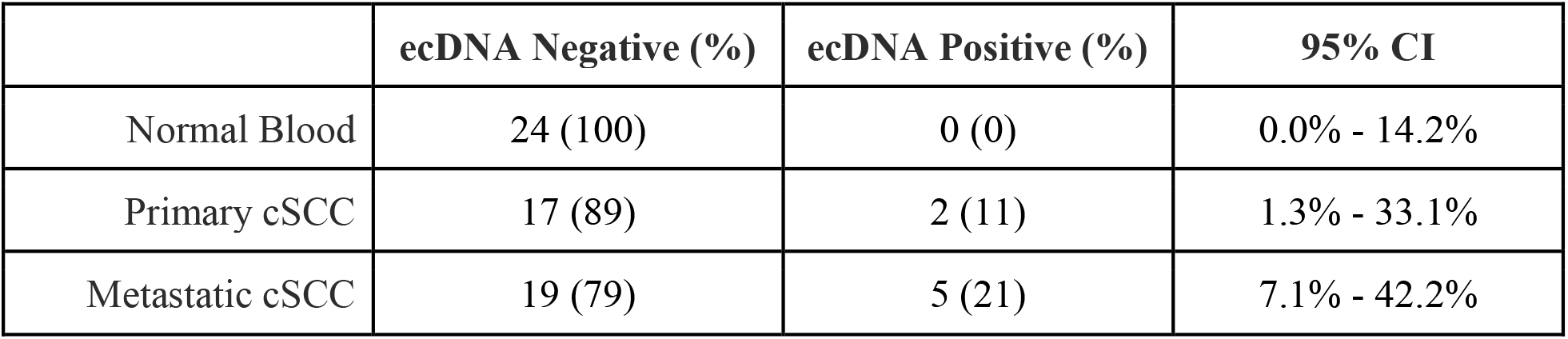
Prevalence of ecDNA in cSCC and normal blood detected by AmpliconArchitect. CI, confidence interval. Confidence intervals computed with binom.test in R.

|  | **ecDNA Negative (%)** | **ecDNA Positive (%)** | **95% CI** |
| --- | --- | --- | --- |
| Normal Blood | 24 (100) | 0 (0) | 0.0% - 14.2% |
| Primary cSCC | 17 (89) | 2 (11) | 1.3% - 33.1% |
| Metastatic cSCC | 19 (79) | 5 (21) | 7.1% - 42.2% |

On average, cSCC-associated ecDNA amplicons were 1.17 Mb (range 0.1 Mb - 3Mb) and contained 14 genes (range 0 - 61). ecDNA in different tumors from the same patient (CSCC_0010) demonstrated shared genes and nearly identical genomic regions, consistent with a common origin and suggesting ecDNA in primary cSCC is maintained during tumor progression and metastasis. Oncogenes recurrently amplified on ecDNA in other cancer types, such as *MYC, JUNB*, and *CALR* were present on cSCC ecDNA amplicons in 60% (3 of 5) of metastases (**Supplementary Table S1, S2**, and **Figure 1a**). Examination of the frequencies at which these genes are amplified on ecDNA in a larger collection of tumors from The Cancer Genome Atlas (TCGA) and the Pan-Cancer Analysis of Whole Genomes (PCAWG) (Kim et al. 2020) showed that our cSCC cohort was less likely than not to detect recurrent ecDNA amplifications due to sampling variability and the relatively small size of our study (**Figure 1b**). Of the two metastases that contained ecDNA free of canonical oncogenes, one contained only amplified noncoding regions and the other had an amplicon that included *RBPJ*, which is upregulated early in cSCC development and promotes tumorigenesis (Al Labban et al.). We next analyzed paired RNA sequencing (RNA-Seq) data from cSCC metastases and demonstrated higher expression of ecDNA amplicon genes in tumors with ecDNA, corroborating earlier findings of enhanced chromatin accessibility and oncogene transcription in ecDNA (Wu et al. 2019) (**Figure 1c**). Taken together, these results indicate that ecDNA in cSCC involves oncogenes known to be recurrently amplified on ecDNA in different cancer types as well as other cancer-promoting genes and confirm an effect on transcription.

**Figure 1.**
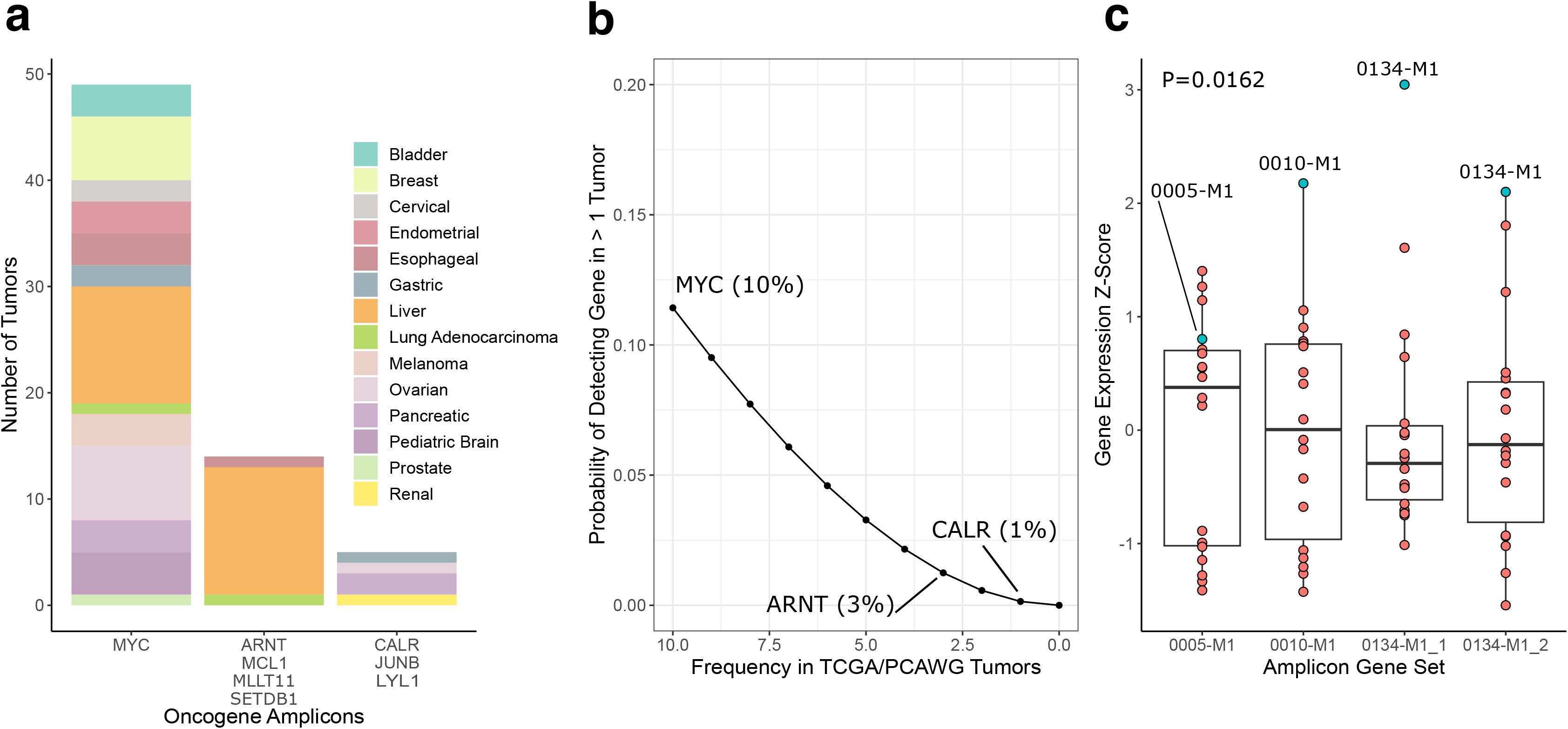
Characteristics of ecDNA in cSCC. (**a**) ecDNA prevalence of oncogenes from cSCC-associated ecDNA in a combined TCGA/PCAWG cohort of 3,212 tumors. Bars represent individual ecDNA amplicons found in cSCC with canonical oncogenes shown. (**b**) Probability of detecting ecDNA oncogene recurrence in our cSCC cohort based on TCGA/PCAWG prevalence. The ecDNA detection frequencies (percent) of *MYC, ARTN*, and *CALR* across 14 cancer types in the TCGA/PCAWG cohort is shown for reference. (**c**) Expression of signatures consisting of genes amplified on cSCC ecDNA were distilled and quantified in 18 metastases that had paired WGS and RNA-Seq data. Each signature consists of the genes found in a specific ecDNA amplicon. Two amplicons were detected in CSCC_0134. Turquoise dots represent the expression score of the tumor that a given signature was derived from. Red dots represent tumors that did not possess that specific ecDNA amplicon. In three of four signatures, the tumor used to generate the corresponding ecDNA signature showed the highest expression. Expression scores were calculated using GSVA. P-value calculated using Welch’s T-test.

Tumors with ecDNA have been associated with worse clinical outcomes (Kim et al. 2020), raising the question of whether the presence of ecDNA in cSCC might similarly herald a poor prognosis. We did not identify correlations between ecDNA status and age, nodal status, extracapsular spread, grade, or immune status, but recognize that with few ecDNA-positive samples, the power of our study to detect associations is low (**Table 2)**. Recent studies in various cancer types including cSCC indicate that the lymph node ratio (LNR), or the number of positive lymph nodes divided by the total nodal yield, can improve prediction of tumor recurrence and overall survival by considering the impact of both positive and normal nodes (Tseros et al. 2016; Vasan et al. 2018). Logistic regression analysis demonstrated a significantly higher LNR in ecDNA-positive cSCC compared to tumors without ecDNA, suggesting ecDNA in cSCC is associated with more aggressive disease (**Table 2** and **Figure 2a**). To determine whether ecDNA might arise in the setting of increased gene mutation, we compared the tumor mutational burdens (TMB) of cSCC with and without ecDNA. A correlation between TMB and ecDNA status was not identified (**Figure 2b**); however, cSCC with ecDNA showed enrichment of the SBS7 mutational signature that is associated with ultraviolet (UV) light exposure (Alexandrov et al. 2013) (**Figure 2c**). These findings together suggest ecDNA in cSCC may arise through repair of UV-induced DNA damage and be an indicator of more extensive disease.

**Table 2.** Clinical associations with ecDNA in metastatic cSCC. Mean values are presented for age and LNR. Fisher’s exact test was used to calculate P-values for nodal status, extracapsular spread, grade, and immune status. Age P-value was calculated using Welch’s T-test. Lymph node ratio P-value was calculated using logistic regression.

| **Characteristic** | **ecDNA Negative** | **ecDNA Positive** | **P-Value** |
| --- | --- | --- | --- |
| Age | 69.5 | 71.6 | 0.618 |
| Nodal Status |  |  | 0.3905 |
| 3b | 1 | 0 |  |
| N1 | 2 | 0 |  |
| N2 | 2 | 2 |  |
| N3 | 14 | 3 |  |
| Extracapsular Spread |  |  | 0.72 |
| No | 4 | 1 |  |
| Yes | 1 | 1 |  |
| Not stated | 14 | 3 |  |
| Grade |  |  | 0.8287 |
| 1 | 1 | 0 |  |
| 2 | 4 | 0 |  |
| 3 | 11 | 4 |  |
| Not stated | 3 | 1 |  |
| Immune Status |  |  | 1 |
| Competent | 17 | 5 |  |
| Suppressed | 2 | 0 |  |
| Lymph Node Ratio | 0.246 | 0.455 | 0.00901 |

**Figure 2.**
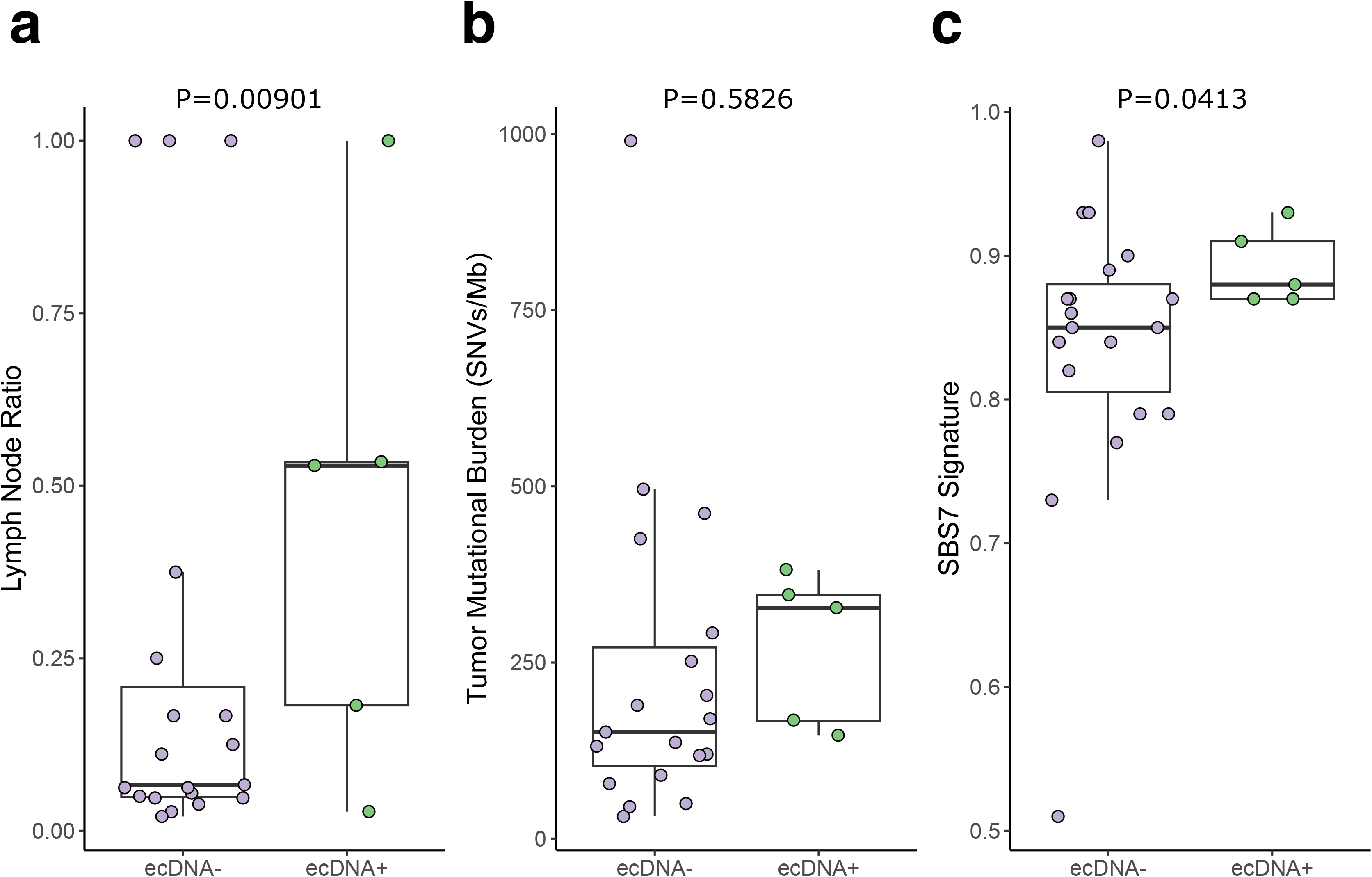
Clinical and molecular associations with ecDNA in metastatic cSCC. (**a**) Lymph node ratio in ecDNA-negative vs. ecDNA-positive tumors. P-value calculated with logistic regression. (**b, c**) Tumor mutational burden and COSMIC SBS7 UV exposure signatures in ecDNA-negative and ecDNA-positive tumors. P-values calculated using Welch’s T-test.

This study expands the spectrum of cancer types harboring ecDNA amplifications to include cSCC, one of the most common human malignancies. cSCC-associated ecDNA possesses features similar to those reported in other cancers and its presence is associated with higher LNRs, suggesting ecDNA status may enable more effective risk stratification of cSCC patients. Our data do not identify a difference in ecDNA prevalence between primary and metastatic cSCC or detect recurrent ecDNA amplifications in cSCC; however, given the relatively small sample size of our study, these results require confirmation in larger cohorts. The presence of ecDNA in primary tumors raises the possibility that extrachromosomal amplifications may arise early in disease progression. A previous report detected ecDNA at higher frequencies and levels in esophageal adenocarcinoma compared to Barrett’s esophagus (Luebeck et al. 2023), suggesting ecDNA may undergo similar positive selection during the progression of premalignant actinic keratoses to early in situ carcinomas and invasive cSCC. Given the association between ecDNA and worse outcomes, another open question is the extent to which ecDNA is present in other cSCC subtypes, such as keratoacanthomas that display benign clinical behavior or aggressive tumors that develop in patients with recessive dystrophic epidermolysis bullosa. Future studies documenting the prevalence and characteristics of ecDNA during cSCC progression or in tumors with distinct clinical phenotypes will help clarify the contribution of ecDNA in this malignancy.

## Data Availability Statement

Code for this analysis is available at https://github.com/tjbencomo/ecdna-cscc. ecDNA amplicon data is available in the supplementary data file. The variant call format files have been deposited at the European Genome-Phenome Archive, which is hosted by the EMBL-European Bioinformatics Institute and the Center for Genomic Regulation, under accession number EGAS00001006378.

The authors declare no potential conflicts of interest.

## Acknowledgements

We thank Vanessa Lopez-Pajares for helpful discussions and also wish to acknowledge the Stanford Research Computing and the National Computational Infrastructure of Australia for providing computational resources that contributed to these results.

## Author Contributions Statement

Conceptualization: TB (lead), AT (supporting), BA (supporting), MR (supporting), CSL (lead); Data curation: TB (equal), AT (equal), BA (equal), MR (equal); Formal analysis: TB (leading); Investigation: TB (lead), AT (supporting), BA (supporting), MR (supporting); CSL (lead); Methodology: TB (lead), CSL (supporting); Visualization: TB (lead), CSL (supporting), Writing - original draft: TB (equal), CSL (equal); Writing - review & editing: TB (lead), AT (supporting), BA (supporting), MR (supporting), CSL (lead); Funding acquisition: BA (equal), MR (equal), CSL (equal).

## Supplementary Materials and Methods

### Sample Collection

Sequencing of metastatic cSCC was performed with Institutional Human Research Ethics approval (UOW/ISLHD HREC14/397) as previously described (Thind et al. 2022). Sequencing of primary cSCC was previously performed with Sydney Local Health District Human Research Ethics approval (X19-282, Gupta et al. 2023). All patients provided written, informed consent.

### Bioinformatic Analysis

Raw FASTQ files were processed using the AmpliconArchitect pipeline to detect ecDNA amplicons (Turner et al. 2017). RNA-Seq reads were aligned using splice-aware aligner STAR (v2.7.2) (Dobin et al. 2013) and quantified with RSEM (Li and Dewey 2011). The count data was transformed with the variance stabilizing transformation (VST) using DESeq2 (DESeq2_1.30.1) (Love et al. 2014). ecDNA amplicon gene signatures were scored using GSVA (Hänzelmann et al. 2013) on VST count data. Somatic mutations, tumor mutational burden, and enrichment of mutational signature scores were distilled from previously generated data (Thind et al. 2022). Probabilities in Figure 1b were calculated using 1 - pbinom(1, 6, Kim 2020 prevalence). The P-value for the difference in lymph node ratio was calculated using an aggregated mixed effects logistic regression model that considered the number of positive and total lymph nodes removed for each patient. A random effect was included for each patient. Statistical calculations and data wrangling were performed with R (v4.3.1).

